# C2-α-dicarbonyls selectively impair morphodynamic interactions between cells and extracellular matrix

**DOI:** 10.1101/2025.07.04.663109

**Authors:** Bidita Samanta, Prosenjit Sen, Ramray Bhat

**Affiliations:** Department of Bioengineering, Indian Institute of Science, Bengaluru, 560012, India; Centre for Nano Science and Engineering, Indian Institute of Science, Bengaluru, 560012, India; Department of Developmental Biology and Genetics, Indian Institute of Science, Bengaluru, 560012, India

**Keywords:** Type 2 Diabetes, Dicarbonyl, Laminin rich basement membrane, Collagen I, migration, adhesion

## Abstract

Ageing and diabetes, risk factors for cardiovascular diseases and cancer, are characterized by abnormally high tissue levels of several dicarbonyl species. Although all dicarbonyls disrupt biomolecular structure and function by forming advanced glycation end products, understanding the relative toxicity of each can help design targeted control strategies. Although known for cells, the comparative effects of distinct dicarbonyls on extracellular matrices (ECM) have yet to be benchmarked. Here, we investigate how glycation by three well known dicarbonyls: the C2-α-dicarbonyls methylglyoxal (MGO) and glyoxal (GO), and the C6-α-dicarbonyl 3-deoxyglucosone (3-DG), affects two ECMs, Collagen I and laminin-rich basement membrane (lrBM), and alters their morphodynamic interactions with two human cells: untransformed endothelial TeloHAEC and aggressive triple negative breast cancer MDA-MB-231. On dicarbonyl-treated lrBM, endothelial and cancer cell adhesion, collective cord morphogenesis, cancer cell motility, shape polarity and its entropy are perturbed at low MGO levels, high GO levels, and are unchanged for 3-DG. Alcian Blue staining of lrBM shows dissolution of matrix at low MGO and high GO levels with no effect on 3-DG exposure. On collagen I substrata, adhesion for both cells is disrupted at low MGO levels, at high GO levels but is unchanged for 3-DG. Interestingly at high concentrations of MGO and GO, the latter decreases motility and increases elongation entropy more than the former; 3-DG has no effect. Picrosirius staining suggests that at high levels GO-driven collagen hyperpolymerization is similar to MGO with no effect of 3-DG. Our observations therefore suggest qualitative and quantitative distinctions in the effect of C2- and C-6-α-dicarbonyls on ECM homoeostasis and ECM-driven cellular morphodynamics.

## INTRODUCTION

Type 2 Diabetes mellitus (T2DM), a chronic metabolic disorder associated with deregulated insulin signaling, is accompanied by an increase in blood glucose levels^1,2^. Hyperglycemia increases interstitial glucose in insulin-independent tissues like red blood cells, peripheral nerve tissue cells, endothelial cells, eye lens cells, and kidney epithelia^3^. It increases metabolic (glycolytic) flux and impairs the dicarbonyl detoxification system resulting in accumulation of dicarbonyl metabolites, also referred to as dicarbonyl stress^4^. Dicarbonyls are electrophiles that react with proteins, peptides, amino acids, phospholipids, and nucleotides and undergo complex chemical rearrangements like oxidation, reduction, and hydration, to form heterogeneous, stable, and irreversible end-products called advanced glycation end products (AGEs)^5–7^. Essential but pathological reactions involved in the formation of AGEs include Maillard reaction (non-enzymatic reaction that occurs between reducing sugars and amino groups)^3,8^, Polyol pathway (metabolic pathway that involves conversion of excess glucose to fructose via sorbitol)^9,10^, oxidation of glucose, and peroxidation of lipids^5,6^. The reaction of dicarbonyls with proteins inside cells alters their structures leading to mitochondrial dysfunction, formation of reactive oxygen species (ROS), apoptosis, anoikis, etc, whereas their reaction with nucleotides leads to DNA strand breaks and mutations^7,11,12^. The accumulation of glycated proteins leads to chronic histopathological complications associated with T2DM including retinopathy, nephropathy, neuropathy, etc^13^. Even newly diagnosed T2DM patients without any other complications have been reported to show a 1.62 fold increase in the presence of methylglyoxal, a well-known dicarbonyl, in the plasma^14,15^.

AGEs can be formed by multiple reactive carbonyls. Examples include diacetyl, glyoxal, methylglyoxal, glyceraldehyde, glycolaldehyde, 1-deoxyglucosone, and 3-deoxyglucosone^6^. Similarity in the crosslinks and receptor recognition factors of AGEs formed by α-oxoaldehydes like methylglyoxal (MGO), glyoxal (GO) and 3-deoxyglucosone (3-DG) with glucose-derived AGEs makes them the most widely studied of glycating agents^13^. These dicarbonyl metabolites have endogenous and exogenous sources. Pathways that result in formation of MGO include anaerobic glycolysis, gluconeogenesis, glyceroneogenesis, ketone body metabolism, monosaccharide degradation, etc. GO is formed by metabolic pathways like glycolysis, lipid peroxidation, glycated protein degradation, oxidative degradation of serine, nucleotide degradation, etc. 3-DG is produced by enzymatic repair of glycated proteins, glycated protein degradation, monosaccharide degradation, fructose metabolism, etc^6,7,16–18^. Exogeneous sources include various food and beverages ^6,7^. While MGO and GO are detoxified with the help of glutathione-dependent glyoxylase, aldoketo-reductases, and aldehyde dehydrogenase, 3-DG is detoxified with the help of aldoketo-reductases and aldehyde dehydrogenase^7^. Pathological conditions like T2DM impair these detoxification pathways, increasing dicarbonyl levels by up to three-fold^6^. MGO is a C2-α-dicarbonyl with a ketone and an aldehyde group, GO is also an C2-α-dicarbonyl but with two aldehyde groups, while 3-DG is a C6-α-dicarbonyl with one ketone and one aldehyde group^19^. Previous studies of these dicarbonyls on transformed keratinocytes reported MGO and GO increase cytotoxicity, reduce cell proliferation and alter cell cycle more than 3-DG^20^. Amongst the three dicarbonyls, MGO has higher reactivity than GO and 3-DG as reported by the higher apoptosis of keratinocytes, higher reactivity with GSH (glutathione), and higher stimulation of VEGF (Vascular endothelial growth factor) production by peritoneal mesenchymal and endothelial cells^19–21^. While MGO and GO are both C2-α-dicarbonyls, they show differences when it comes to reactivity, indicating the context-dependent roles of the dicarbonyls. On one hand, MGO interacts preferentially with arginine but also with lysine and cysteine residues, GO reacts preferentially with lysine but does react with arginine and cysteine too^22,23^. 3-DG first reacts with arginine groups in a protein followed by lysine residues^24^. This brings us to the impact of all these dicarbonyls on extracellular matrix proteins like collagen, laminin, fibronectin, elastin, consisting of these amino acids. There are reports on increased crosslinking of Collagen I, dissolution of laminin-rich basement membrane, destabilization of Collagen IV, decreased elasticity of elastin, and increased stiffness of collagen with the administration of dicarbonyl stressors, especially with MGO and GO^22,25,26^. However, a systemic comparison of the three most common dicarbonyl stressors (MGO, GO and 3-DG) on ECM and their effect on physiological and pathological processes is yet to be addressed.

Our work compares the impact on adhesion and migration of two cell lines on two different types of extracellular matrices treated with three different dicarbonyls. Our experiments include a non-cancerous endothelial cell line TeloHAEC and a triple breast cancer cell line MDA-MB-231. This allows a broad spectrum of experimentation since T2DM is known to cause endothelial dysfunction and increase the risk of developing cancers^18,27^. The ECMs used in this study include Collagen I and Laminin-rich Basement membrane lrBM. Both collagen and lrBM allow physiologically relevant cell matrix interactions due to the presence of cell binding epitopes^28^. The three dicarbonyls that have been used to mimic diabetic microenvironment in the experiments we perform include MGO, GO and 3-DG. They are the first reactive carbonyls to be identified as benzo-quinoxaline derivatives using HPLC with 2,3-diaminonaphthalene (DAN) reagent^29^. MGO is the most extensively used model for studying dicarbonyl stress due to its high reactivity and highest endogenous flux^11,15^. GO and 3-DG are also markers of chronic hyperglycemic and diabetic complications^30^. Our results serve to show how the C2-α-dicarbonyl exert far stronger matritoxic effects perturbing cellular behavior.

## RESULTS

### Adhesion and migration of endothelial cells on LrBM are perturbed by exposure to MGO and GO, but not 3-DG

Increase in dicarbonyl stress, especially due to MGO, is known to cause vascular complications and endothelial dysfunction; hence, the human vasculature represents a pertinent system to compare the relative toxicity of the dicarbonyls^18^. Laminin-rich basement membrane (lrBM) closely mimics the mechanical and chemical properties of the vascular extracellular matrix^31^. We sought to investigate the impact of C2-α-dicarbonyls methylglyoxal (MGO) and glyoxal (GO), and the C6-α-dicarbonyl 3-deoxyglucosone (3-DG) on the adhesion and migration on lrBM of endothelial cells (TeloHAEC). Matrix adhesion is critical for untransformed cells like endothelia to form a monolayer within lumen-containing organs^32^. Adhesion is also a necessary step towards migration on such matrices, which along with cell division contributes to physiological ordering of such homoeostatic monolayers^33^.

Adhesion of TeloHAECs on lrBM was studied by adding cells to lrBM, previously treated with MGO, GO and 3-DG for 24 h (schematic of experimental assay shown in Figure 1Ai). To confirm that the cells only interact with the treated matrix and not with residual dicarbonyl stressors, lrBM was properly washed with 1X PBS. The dicarbonyl stressors before addition on lrBM and the solution collected after washing lrBM with 1X PBS were allowed to react with chromic acid. Chromic acid changes from orange to blue in presence of aldehydes; reaction of chromic acid with dicarbonyls before addition gave blue color and after wash gave orange color, indicating the absence of dicarbonyl stressors after the wash (Supplementary Figure 1A). Unlike untreated control conditions, where cells attached to lrBM, increasing concentrations of 50 μM, 100 μM and 200 μM MGO significantly decreased the number of cells adhered to the matrix (Figure 1Aii top row, p<0.0001 for 50 μM, 100 μM and 200 μM of MGO-treated lrBM significance tested using one way ANOVA with Dunnett’s multiple comparisons; data collected for three biological replicates with 10 cells for each replicate). GO shows negligible attachment of cells to the matrix only at the highest concentration of 200 μM (Figure 1Aii middle row, ns for 50 μM and 100 μM and p<0.0001 for 200 μM of GO-treated lrBM; significance tested using one way ANOVA with Dunnett’s multiple comparisons; data collected for three biological replicates with 10 cells for each replicate). 3-DG did not perturb the adhesion of endothelial cells to lrBM and no significant differences could be observed in the adhesion of the cells when compared with untreated conditions (Figure 1Aii bottom row, ns for 50 μM, 100 μM and 200 μM of 3-DG-treated lrBM; significance tested using one way ANOVA with Dunnett’s multiple comparisons; data collected for three biological replicates with 10 cells for each replicate).

**Figure 1.**
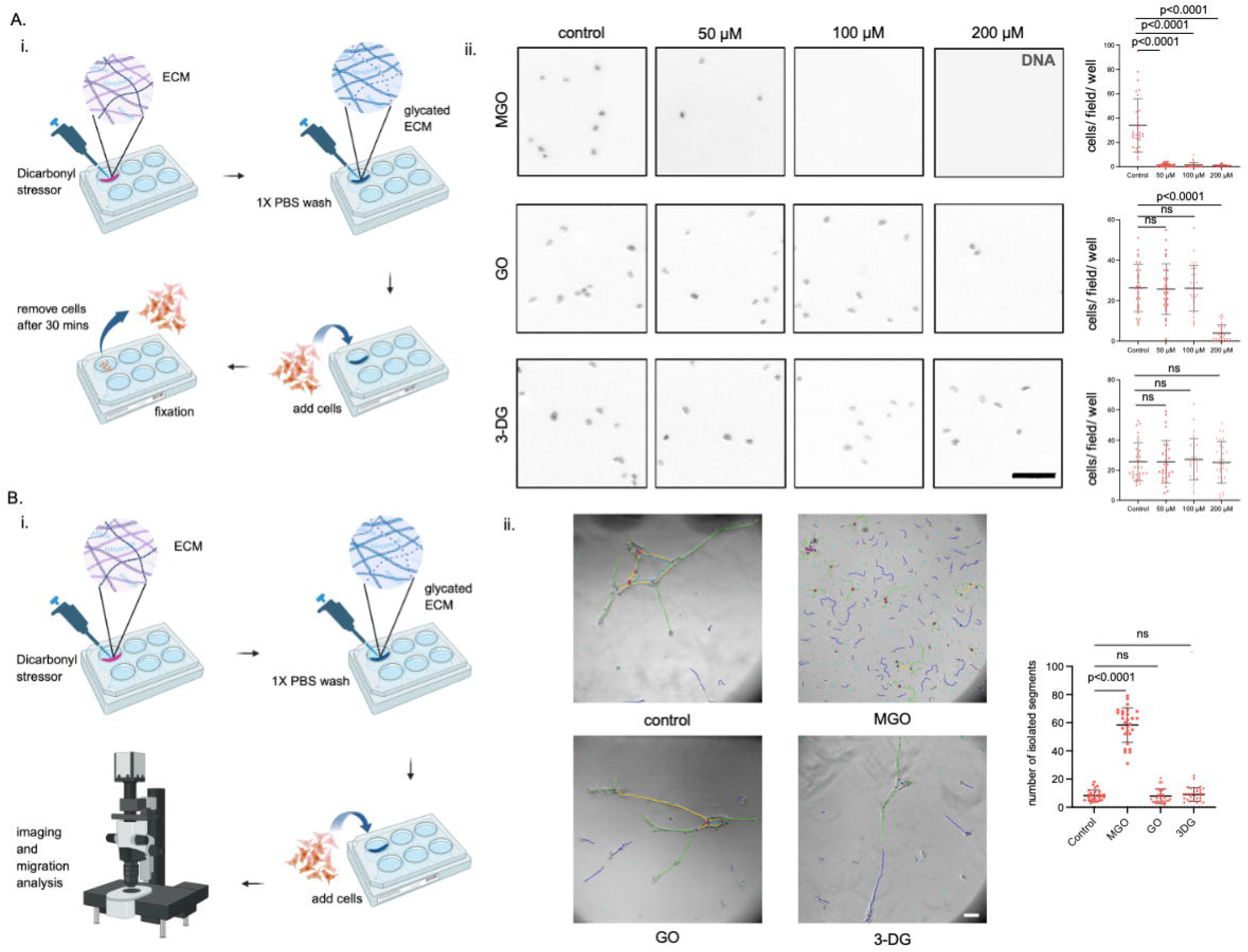
Higher exposure to MGO perturbs the adhesion and migration of endothelial cells on lrBM to a greater extent than GO and 3-DG. (Ai) Schematic depiction of the protocol used for cell adhesion assay (Aii) DAPI (black) stained fluorescent images for endothelial cells TeloHAEC adhered to lrBM coats, untreated (left) and treated with 50 μM, 100 μM and 200 μM of methylglyoxal (top), glyoxal (middle) and 3-deoxyglucosone (bottom) for 1 d and scatter plot graphs showing number of cells attached per field on lrBM coats for endothelial cells TeloHAEC, untreated (leftmost) and treated with increasing concentrations of 50 μM, 100 μM and 200 μM of methylglyoxal (top, left to right), of glyoxal (middle, left to right) and of 3-deoxyglucosone (bottom, left to right) for 1 d. (Bi) Schematic depiction of the protocol used for cell migration assay (Bii) Brightfield images of endothelial cells TeloHAEC with meshed pseudo-capillary network structure indicating endothelial tube formation on attachment to lrBM, untreated (left top) and treated with 50 μM of methylglyoxal (right top), 50 μM of glyoxal (left bottom) and 50 μM of 3-deoxyglucosone (right bottom) for 1 d and scatter plot graph (rightmost) of the number of isolated elements formed in the pseudo capillary network structure of endothelial cells TeloHAEC, untreated and treated with 50 μM of methylglyoxal, 50 μM of glyoxal and 50 μM of 3-deoxyglucosone (left to right) for 1 d. Significance for (A) tested using one way ANOVA with Dunnett’s multiple comparisons; data collected for three biological replicates with 10 cells for each replicate. Significance for (B) tested using one way ANOVA with Dunnett’s multiple comparisons; data collected for three biological replicates with 28 fields in total. Error bars represent mean ± SD. Scale Bar for (A) and (B) = 100 μM. Schematics Ai and Bi are prepared in biorender. Three or more independent repeats were performed for each experiment.

The data obtained from adhesion studies indicates a change in the cellular properties at 50 μM of MGO. This prompted us to study the migration of TeloHAEC after treating lrBM with 50 μM of MGO, GO and 3-DG for 1 d. Cells were added to the matrix after 1 d of treatment and migration was analyzed after 20 h of incubation (schematic of experimental assay shown in Figure 1Bi). Endothelial cells tend to form meshed pseudo-capillary network structure on lrBM. Laminin, the primary component of lrBM, leads to “vascular mimicry” by endothelial cells that can initiate capillary-like multicellular tubular structures^34,35^. Hence, under untreated conditions, TeloHAEC were observed to form meshed networks (Figure 1Bii left top untreated control). On lrBM-treated MGO at 50 μM, meshed networks did not form but rather remained as isolated cells or clusters; in contrast, they did form on lrBM treated with equivalent concentrations of GO and 3-DG (Figure 1Bii middle top, left bottom, middle bottom). This is quantified in the form of a graph of number of isolated segments (Figure 1Bii right). The number of isolated segments is higher in case of MGO, but very low in, and insignificantly different between, untreated controls and GO-and 3-DG-treated conditions (Figure 1Bii Number of isolated segments, p<0.0001 for MGO, ns for GO and ns for 3-DG; significance tested using one way ANOVA with Dunnett’s multiple comparisons; data collected for three biological replicates with 28 fields in total).

These experiments indicated that GO tends to interact with ECM in a manner like MGO at higher concentrations, where it allows negligible cell adherence at 200 μM and like 3-DG at lower concentrations, where it allows cells to attach at 50 μM and 100 μM. This is also observed in cell migration observed at the concentration of 50 μM where GO tends to form meshes similar to 3-DG and untreated samples.

### Adhesion and migration of cancer cells on LrBM are perturbed by exposure to MGO and GO, but not 3-DG

We next sought to investigate how glycation alters the laminin-rich matrix milieu around cancer cells and whether the same affects their behavior. This was studied by adding cells to lrBM, previously treated with MGO, GO and 3-DG for one d. While untreated conditions allowed the triple negative breast cancer MDA-MB-231 cells to strongly adhere to lrBM, treating the matrix with increasing concentrations of 50 μM, 100 μM and 200 μM MGO attenuated cell adhesion progressively to greater extents. (Figure 2A top row, p<0.0001 for 50 μM, 100 μM and 200 μM of MGO-treated lrBM; significance tested using one way ANOVA with Dunnett’s multiple comparisons; data collected for three biological replicates with 10 cells for each replicate). GO shows negligible attachment of cells to the matrix only at the highest used concentration of 200 μM (Figure 2A middle row, ns for 50 μM and 100 μM and p<0.0001 for 200 μM of GO-treated; significance tested using one way ANOVA with Dunnett’s multiple comparisons; data collected for three biological replicates with 10 cells for each replicate). No significant differences can be observed in the adhesion of the cancer cells to 3-DG treated lrBM when compared against untreated controls (Figure 2A bottom row, ns for 50 μM, 100 μM and 200 μM of 3-DG-treated lrBM; significance tested using one way ANOVA with Dunnett’s multiple comparisons; data collected for three biological replicates with 10 cells for each replicate).

**Figure 2.**
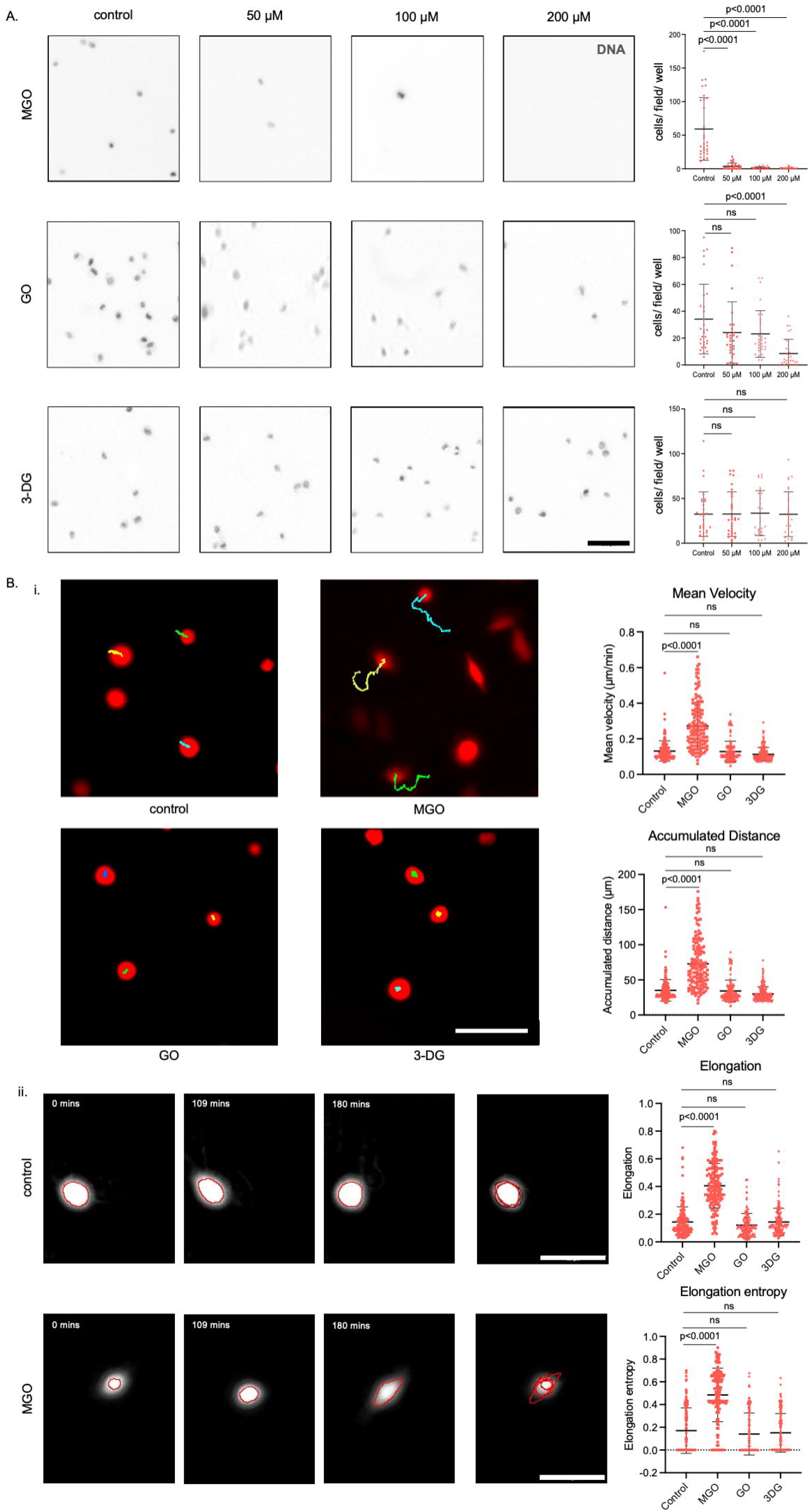
Higher exposure to MGO perturbs the adhesion and migration of cancer cells on lrBM to a greater extent than GO and 3-DG. (A) DAPI (black) stained fluorescent images for cancer cells MDA-MB-231 adhered to lrBM coats, untreated (left) and treated with 50 μM, 100 μM and 200 μM of methylglyoxal (top), glyoxal (middle) and 3-deoxyglucosone (bottom) for 1 d and scatter plot graphs showing number of cells attached per field on lrBM coats for cancer cells MDA-MB-231, untreated (leftmost) and treated with increasing concentrations of 50 μM, 100 μM and 200 μM of methylglyoxal (top, left to right), of glyoxal (middle, left to right) and of 3-deoxyglucosone (bottom, left to right) for 1 d. (Bi) Red fluorescent images of RFP labelled cancerous cells MDA-MB-231 (left) with their migration tracks for 3 h after attachment to lrBM, untreated (top left) and treated with 50 μM of methylglyoxal (top middle), 50 μM of glyoxal (bottom left) and 50 μM of 3-deoxyglucosone (bottom middle) for 1 d and scatter plot graphs (right) of the mean velocity (right top) and accumulated distance (right bottom) of cancerous cells MDA-MB-231, untreated and treated with 50 μM of methylglyoxal, 50 μM of glyoxal and 50 μM of 3-deoxyglucosone (left to right) for 1 d (Bii) White fluorescent images with red outlines marking the elongation of MDA-MB-231 cells on untreated control lrBM (left top) and on lrBM treated with 50 μM of methylglyoxal (left bottom) at 0 min, 109 min and 180 min (left to right), their change in elongation dynamics (middle) and the scatter plot graphs (right) of elongation (right top) and elongation entropy (right bottom) of cancerous cells MDA-MB-231, untreated and treated with 50 μM of methylglyoxal, 50 μM of glyoxal and 50 μM of 3-deoxyglucosone (left to right) for 1 d. Significance for (A) tested using one way ANOVA with Dunnett’s multiple comparisons; data collected for three biological replicates with 10 cells for each replicate. Significance for (B) tested using one way ANOVA with Dunnett’s multiple comparisons; data collected for three biological replicates with 50 cells for each replicate. Error bars represent mean ± SD. Scale Bar for (A) and (B) = 100 μM. Three or more independent repeats were performed for each experiment.

Cancer cells also show a significant change at the lowest concentration of 50 μM in case of MGO treatment like endothelial cells. Hence, we assayed migration of cancer cells on lrBM treated with 50 μM of dicarbonyls after 1 d of treatment. Migration was analyzed after 20 h of incubation.

MDA-MB-231 on untreated lrBM substrata were rounded in shape; treatment of lrBM with 50 μM of MGO led to cell assuming spindle shape which phenocopies shape on plastic or on glass. In contrast, cells remained circular on GO- and 3-DG-treated lrBM. This was better quantified by measuring their elongation dynamics, which is significantly higher for MGO, but not in case of GO and 3-DG, when compared against untreated conditions (Figure 2Bii Elongation, p<0.0001 for MGO, ns for GO, ns for 3-DG; significance tested using one way ANOVA with Dunnett’s multiple comparisons; data collected for three biological replicates with 50 cells for each replicate)^36^. The cells on untreated, GO- and 3-DG-treated matrix remain circular with time while cells on MGO-treated matrix become more elongated with time (Figure 2Bii Elongation entropy, p<0.0001 for MGO, ns for GO, ns for 3-DG; significance tested using one way ANOVA with Dunnett’s multiple comparisons; data collected for three biological replicates with 50 cells for each replicate). Spindle-shaped cells allow faster motion and cover greater distances, when compared to circular cells, which is evident from the results obtained for mean velocity and accumulated distance, both of which show significant increase on MGO-treated lrBM, but no significant difference in case of GO and 3-DG, when compared against untreated control matrix (Figure 2Bi Mean velocity, p<0.0001 for MGO, ns for GO, ns for 3-DG; Accumulated distance, p<0.0001 for MGO, ns for GO, ns for 3-DG; significance tested using one way ANOVA with Dunnett’s multiple comparisons; data collected for three biological replicates with 50 cells for each replicate).

Our experiments with MDA-MB-231 were consistent with those using TeloHAEC in that treatment with GO is comparable to treatment with 3-DG at lower concentrations, while it is comparable to treatment with MGO at higher concentrations.

### MGO and GO, but not 3-DG, leads to dissolution of LrBM

The previous results reflect an impact induced by dicarbonyls on lrBM which we uncover by examining the treated matrices in the absence of cells. Alcian blue stain, that can stain proteoglycans and glycoproteins, stained lrBM untreated control and lrBM treated with increasing concentrations of MGO, GO and 3-DG. LrBM dissolved in the presence of MGO as well as GO, but not 3-DG (Figure 3A and 3B). To negate the possibility of dissolution of lrBM due to change in pH of the dicarbonyls, we look at the pH values of MGO, GO and 3-DG at 50 μM, 100 μM and 200 μM. All the solutions gave a pH value of approximately 7, closer to the physiological pH (Supplementary Figure 1B). Dissolution was more profound for MGO compared to GO where even the lowest concentration 50 μM of MGO leaves behind a plastic substratum. The plastic substrata did not allow cells to adhere to it, leading to negligible adhesion of cells for MGO at lower concentrations and GO at only the highest concentration of 200 μM. Dissolution of matrix confirms the observation of spindle shaped cancer cells for GO-treated and 3-DG-treated lrBM but circular cells for MGO-treated lrBM and more isolated segments in case of MGO-than GO- and 3-DG-treated lrBM.

**Figure 3.**
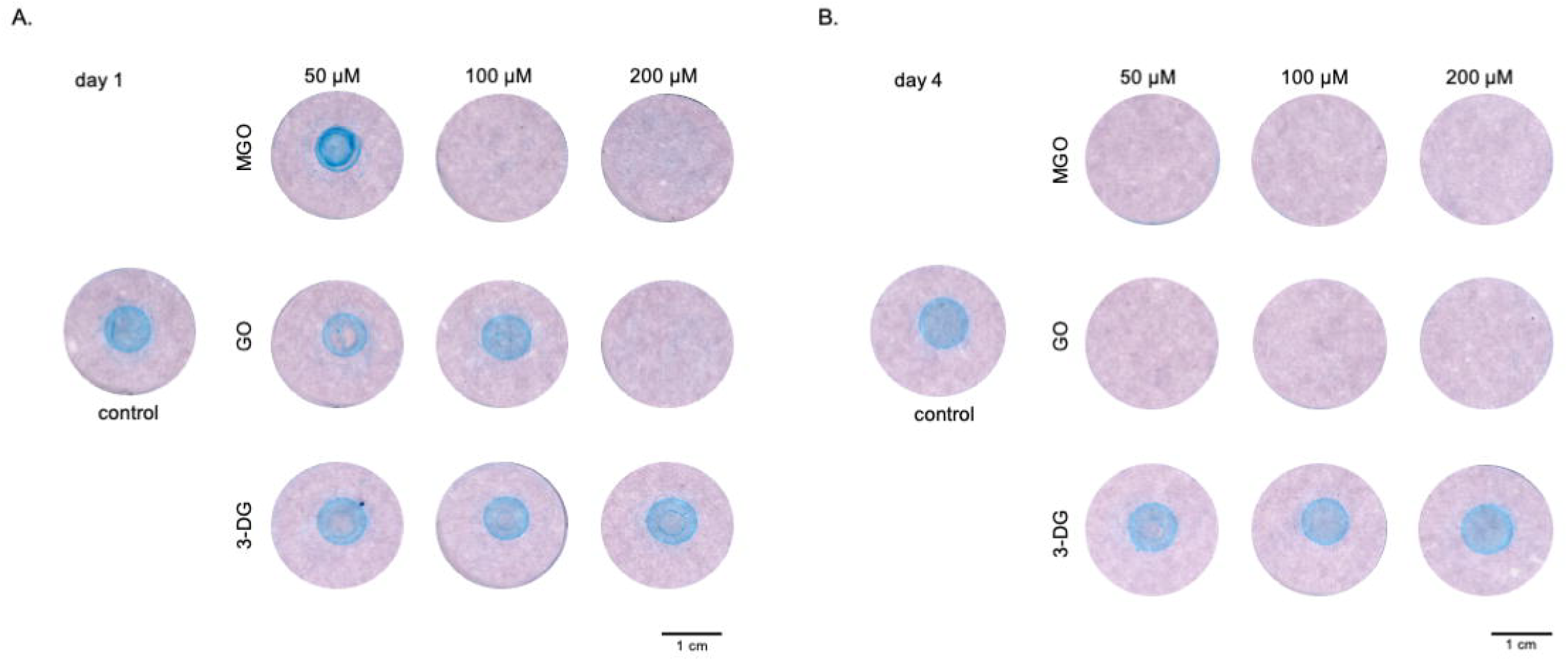
Higher exposure to MGO and GO, but not 3-DG, leads to decreased alcian blue staining of Laminin-rich basement membrane. A) Photomicrographs of solidified lrBM gels stained with alcian blue stain (blue color), untreated (leftmost) and treated with increasing concentration of 50 μM, 100 μM and 200 μM (left to right) of methylglyoxal (top row), glyoxal (middle row) and 3-deoxyglucosone (bottom row) for 1 d. (B) Photomicrographs of solidified lrBM gels stained with alcian blue stain (blue color), untreated (leftmost) and treated with increasing concentration of 50 μM, 100 μM and 200 μM (left to right) of methylglyoxal (top row), glyoxal (middle row) and 3-deoxyglucosone (bottom row) for 4 d. Experiment carried out for more than three biological replicates. Scale bar for (A) and (B) = 1 cm. Three or more independent repeats were performed for each experiment.

### Adhesion and migration of endothelial cells on Collagen I are perturbed by exposure to MGO and GO, but not 3-DG

Dicarbonyl stress leads to diabetic complications in multiple tissues like insulin resistance in skeletal muscles, failure in pancreatic islet function, neurodegeneration, etc^15^. We attempt to understand the impact of dicarbonyl stress on the tissue microenvironment by recruiting Collagen I, most common constituent of the stroma. We examine the effect of glycated Collagen I on the adhesion and migration of endothelial cells.

Adhesion of endothelial cells TeloHAEC on Collagen I was investigated by addition of the cells on Collagen I, previously treated with MGO, GO and 3-DG for 24 h. The number of cells that were able to adhere to Collagen I was observed to be significantly decreasing with increasing concentration and duration of treatment with both MGO and GO. The adhesion of cells to MGO-treated Collagen I significantly reduced, even at the lowest concentration of 50 μM (Figure 4A top row, p<0.0001 for 50 μM, 100 μM and 200 μM of MGO-treated Collagen I; significance tested using one way ANOVA with Dunnett’s multiple comparisons; data collected for three biological replicates with 10 cells for each replicate). This difference was less significant with GO when compared with MGO, with effect observed only at the highest concentration of 200 μM (Figure 4A middle row, ns for 50 μM and 100 μM and p<0.0001 for 200 μM of GO-treated Collagen I; significance tested using one way ANOVA with Dunnett’s multiple comparisons; data collected for three biological replicates with 10 cells for each replicate). No significant difference was observed for 3-DG (Figure 4A bottom row, ns for 50 μM, 100 μM and 200 μM of 3-DG-treated Collagen I; significance tested using one way ANOVA with Dunnett’s multiple comparisons; data collected for three biological replicates with 10 cells for each replicate). This suggests certain structural changes on Collagen I matrix that makes GO-treated matrices react like MGO-treated matrices at higher concentrations and 3-DG-treated matrices at lower concentrations.

**Figure 4.**
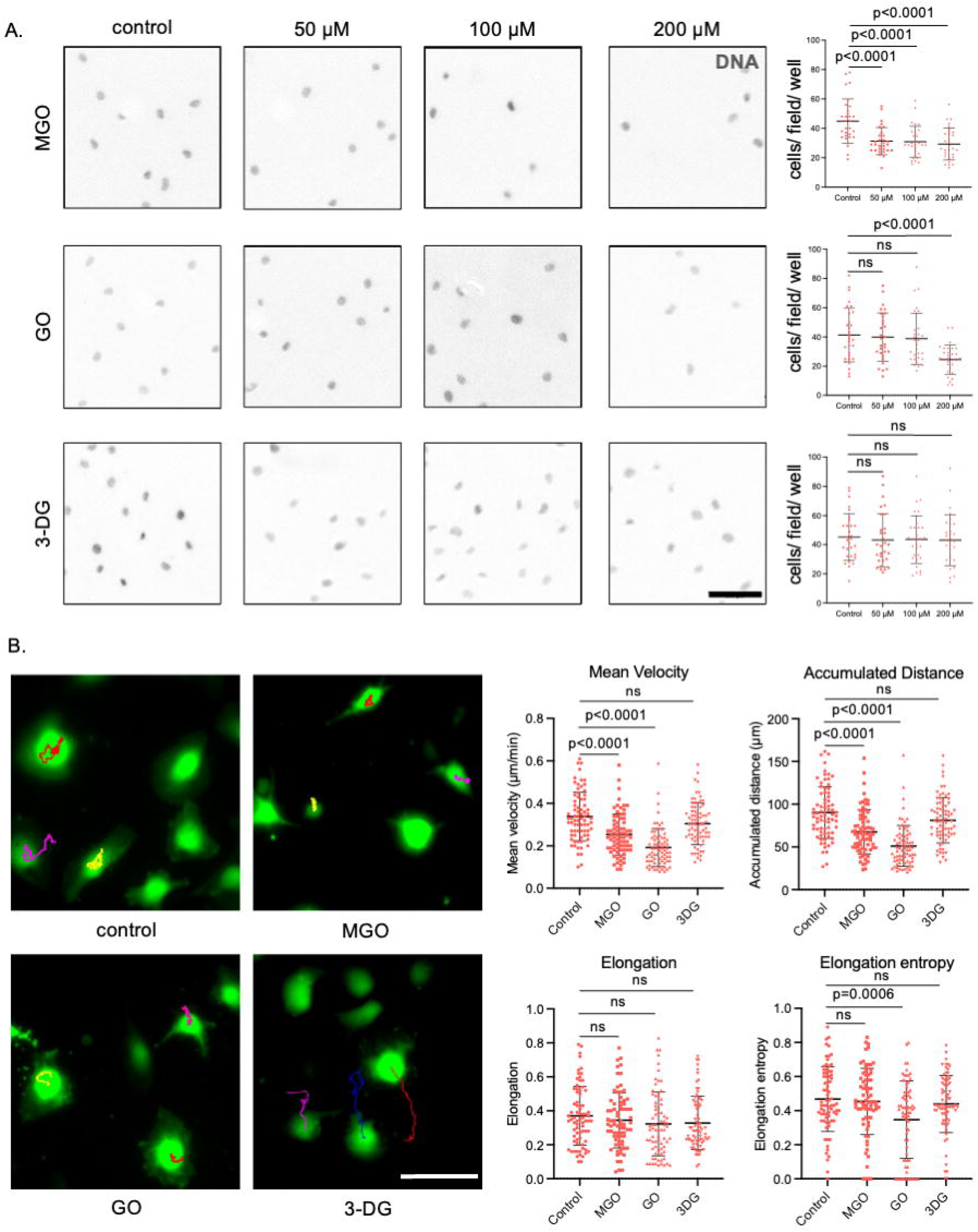
Higher exposure to MGO and GO, but not 3-DG, leads to decreased adhesion and perturbed migration of endothelial cells on Collagen I. (A) DAPI (black) stained fluorescent images for endothelial cells TeloHAEC adhered to Collagen I coats, untreated (left) and treated with 50 μM, 100 μM and 200 μM of methylglyoxal (top), glyoxal (middle) and 3-deoxyglucosone (bottom) for 4 d and scatter plot graphs showing number of cells attached per field on Collagen I coats for endothelial cells TeloHAEC, untreated (leftmost) and treated with increasing concentrations of 50 μM, 100 μM and 200 μM of methylglyoxal (top, left to right), of glyoxal (middle, left to right) and of 3-deoxyglucosone (bottom, left to right) for 4 d. (B) Red fluorescent images of GFP labelled endothelial cells TeloHAEC (left) with their migration tracks for 3 h after attachment to Collagen I, untreated (top left) and treated with 200 μM of methylglyoxal (top right), 200 μM of glyoxal (bottom left) and 200 μM of 3-deoxyglucosone (bottom right) for 4 d and scatter plot graphs (right) of the mean velocity (left top), accumulated distance (right top), elongation (left bottom) and elongation entropy (right bottom) of endothelial cells TeloHAEC, untreated and treated with 200 μM of methylglyoxal, 200 μM of glyoxal and 200 μM of 3-deoxyglucosone (left to right) for 4 d. Significance for (A) tested using one way ANOVA with Dunnett’s multiple comparisons; data collected for three biological replicates with 10 cells for each replicate. Significance for (B) tested using one way ANOVA with Dunnett’s multiple comparisons; data collected for three biological replicates with 25 cells for each replicate. Error bars represent mean ± SD. Scale Bar for (A) and (B) = 100 μM. Three or more independent repeats were performed for each experiment.

Endothelial cells show significant variation in cell adhesion at the concentration of 200 μM of GO. This prompted us to examine migration of endothelial cells at 200 μM concentration of MGO, GO and 3-DG.

Migration of endothelial cells on treated Collagen I was studied after 4 d of treatment with 200 μM of MGO, GO and 3-DG to mimic a stage of vasculopathy with prolonged accumulation of glycating agents. Endothelial cell TeloHAEC assumed spindle shape under untreated control conditions and on treatment with MGO, GO and 3-DG (Figure 4B Elongation, ns for MGO, GO and 3-DG significance tested using one way ANOVA with Dunnett’s multiple comparisons; data collected for three biological replicates with 25 cells for each replicate), but the velocity and accumulated distance decreased for MGO- and GO-treated Collagen I, but not for 3-DG-treated Collagen I (Figure 4B Mean velocity, p<0.0001 for MGO, p<0.0001 for GO and ns for 3-DG; Accumulated distance, p<0.0001 for MGO, p<0.0001 for GO and ns for 3-DG; significance tested using one way ANOVA with Dunnett’s multiple comparisons; data collected for three biological replicates with 25 cells for each replicate).

The results obtained from adhesion and migration of endothelial cells point towards the similarity of GO-treated matrices with MGO-treated matrices at higher concentration and duration of treatment and with 3-DG at lower concentration and duration of treatment. While the number of cells that adhere to GO-treated Collagen I is equivalent to 3-DG-treated Collagen I at 50 μM and 100 μM, it is equivalent to MGO-treated Collagen I at 200 μM. Cell migratory parameters for GO-treated matrix are also like MGO-treated matrix at the measured concentration of 200 μM.

### Adhesion and migration of cancer cells on Collagen I are perturbed by exposure to MGO and GO, but not 3-DG

We next examine how the same glycated microenvironment alters the behavior of cancer cells. Adhesion of breast cancer cells MDA-MB-231 was studied on Collagen I treated with MGO, GO and 3-DG for 24 h. Less number of cells attached to Collagen I with increasing concentration and duration of MGO and GO treatment. The adhesion of cells to MGO-treated Collagen I significantly reduced at all concentrations (Figure 5A top row, p<0.0001 for 50 μM, 100 μM and 200 μM of MGO-treated Collagen I; significance tested using one way ANOVA with Dunnett’s multiple comparisons; data collected for three biological replicates with 10 cells for each replicate). GO showed a significant effect only at the highest concentration of 200 μM (Figure 5A middle row, ns for 50 μM and 100 μM and p=0.0040 for 200 μM of GO-treated Collagen I; significance tested using one way ANOVA with Dunnett’s multiple comparisons; data collected for three biological replicates with 10 cells for each replicate). No change was observed for 3-DG-treated Collagen I (Figure 5A bottom row, ns for 50 μM, 100 μM and 200 μM of 3-DG-treated Collagen I; significance tested using one way ANOVA with Dunnett’s multiple comparisons; data collected for three biological replicates with 10 cells for each replicate).

**Figure 5.**
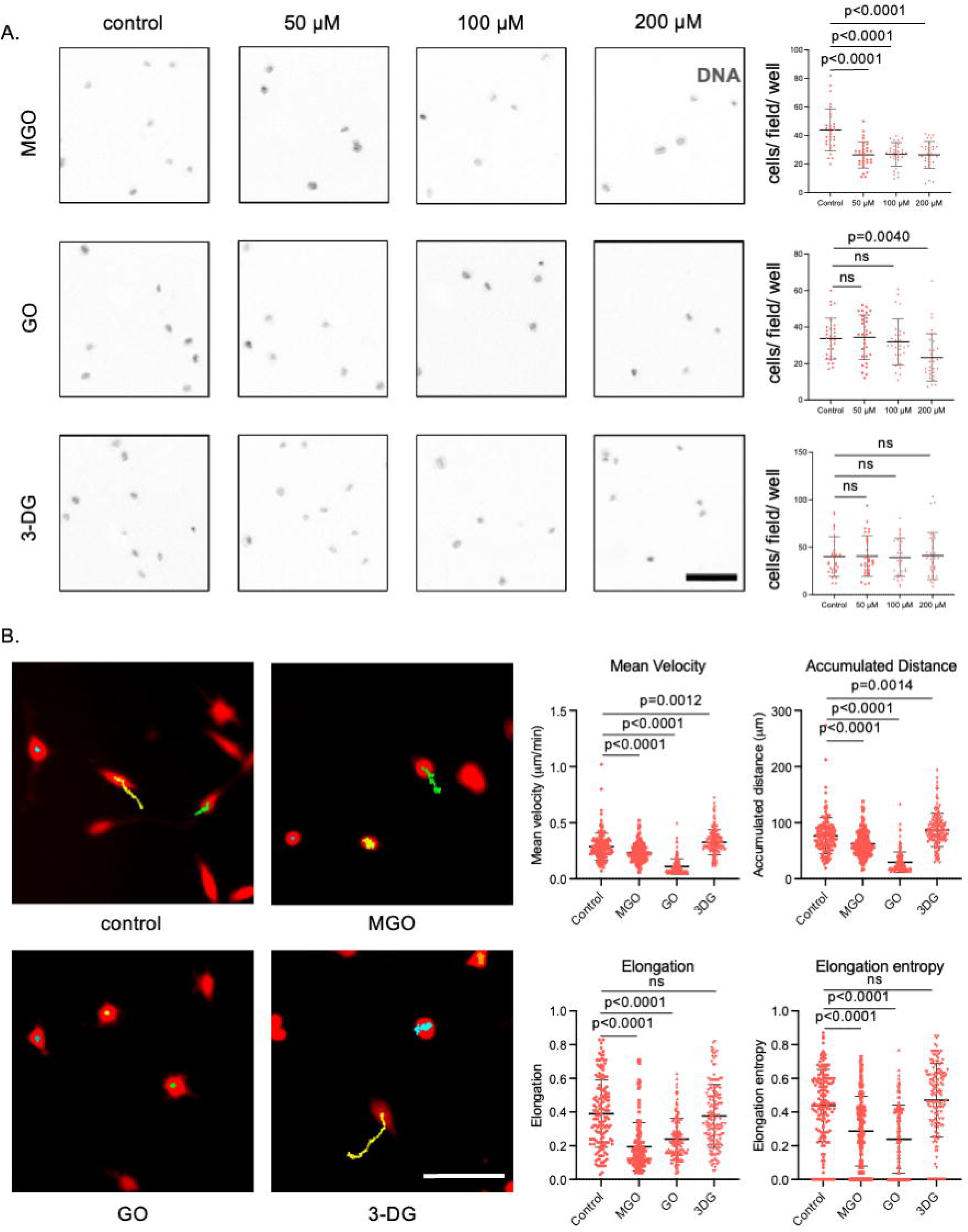
Higher exposure to MGO and GO, but not 3-DG, leads to decreased adhesion and perturbed migration of cancer cells on Collagen I. (A) DAPI (black) stained fluorescent images for cancer cells MDA-MB-231 adhered to Collagen I coats, untreated (left) and treated with 50 μM, 100 μM and 200 μM of methylglyoxal (top), glyoxal (middle) and 3-deoxyglucosone (bottom) for 4 d and scatter plot graphs showing number of cells attached per field on Collagen I coats for cancer cells MDA-MB-231, untreated (leftmost) and treated with increasing concentrations of 50 μM, 100 μM and 200 μM of methylglyoxal (top, left to right), of glyoxal (middle, left to right) and of 3-deoxyglucosone (bottom, left to right) for 4 d. (B) Red fluorescent images of RFP labelled cancerous cells MDA-MB-231 (left) with their migration tracks for 3 h after attachment to Collagen I, untreated (top left) and treated with 200 μM of methylglyoxal (top right), 200 μM of glyoxal (bottom left) and 200 μM of 3-deoxyglucosone (bottom right) for 4 d and scatter plot graphs (right) of the mean velocity (left top), accumulated distance (right top), elongation (left bottom) and elongation entropy (right bottom) of cancerous cells MDA-MB-231, untreated and treated with 200 μM of Methylglyoxal, 200 μM of Glyoxal and 200 μM of 3-Deoxyglucosone (left to right) for 4 d. Significance for (A) tested using one way ANOVA with Dunnett’s multiple comparisons; data collected for three biological replicates with 10 cells for each replicate. Significance for (B) tested using one way ANOVA with Dunnett’s multiple comparisons; data collected for three biological replicates with 50 cells for each replicate. Error bars represent mean ± SD. Scale Bar for (A) and (B) = 100 μM. Three or more independent repeats were performed for each experiment.

Cancer cells like endothelial cells show significant variation in cell adhesion at the concentration of 200 μM. Hence, we studied migration at 200 μM after 4 d of treatment.

MDA-MB-231 showed circular morphology in presence of MGO and GO while they were elongated spindle shaped in untreated control conditions and on treatment with 3-DG (Figure 5B Elongation, p<0.0001 for MGO, p<0.0001 for GO, ns for 3-DG; significance tested using one way ANOVA with Dunnett’s multiple comparisons; data collected for three biological replicates with 50 cells for each replicate). More significant decrease in velocity and accumulated distance was observed for MGO and GO, but less significant in case of 3-DG (Figure 5B Mean velocity, p<0.0001 for MGO, p<0.0001 for GO, p=0.0012 for 3-DG; Accumulated distance, p<0.0001 for MGO, p<0.0001 for GO, p=0.0014 for 3-DG; significance tested using one way ANOVA with Dunnett’s multiple comparisons; data collected for three biological replicates with 50 cells for each replicate).

Cancer cells show consistent results with endothelial cells in that they indicate the similarity of GO-treated matrices with MGO-treated matrices at higher concentration and duration of treatment and with 3-DG at lower concentration and duration of treatment.

### MGO and GO, but not 3-DG, increases crosslinking of Collagen I

While the migration and adhesion studies indicate structural aberrations in Collagen I in the presence of MGO and GO, we ascertain it with the help of an acellular model. Picrosirius staining was used to determine the structural complexities seen in polymerized collagen I gels with increasing concentration (50 μM, 100 μM and 200 μM) and duration (1 d and 4 d) of MGO, GO and 3-DG. Picrosirius has anionic sulphonic groups that attach with basic amino acids in collagen I to produce a pink color stain^37^. The decrease in intensity of stain for MGO and GO suggested decreased availability of basic amino acids for Picrosirius stain to bind to and hence increased crosslinking of Collagen I. 3-DG did not show any alteration in Collagen I as evident from the unchanged intensity of stain with increasing concentration and duration of treatment (Figure 6A and 6B).

**Figure 6.**
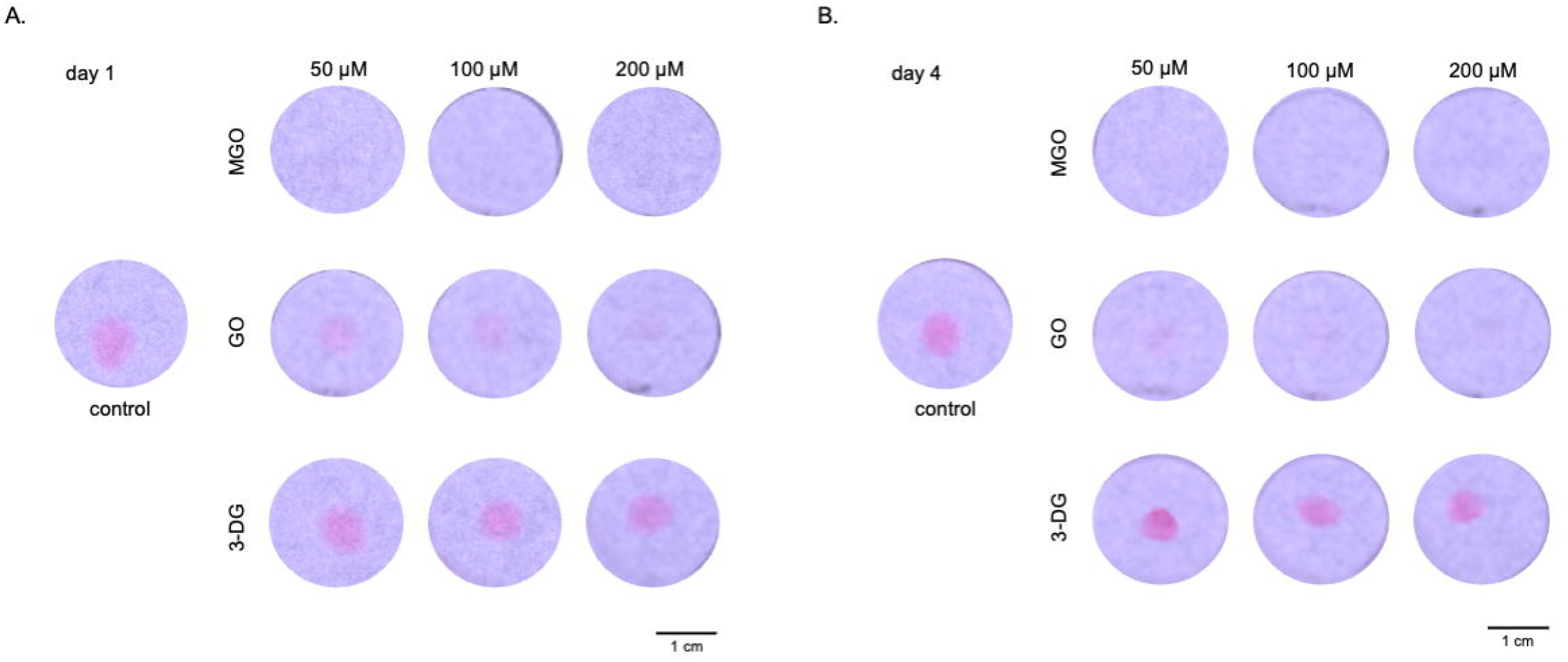
Higher exposure to MGO and GO, but not 3-DG, leads to decreased picrosirius staining of Collagen I. (A) Photomicrographs of polymerized Collagen I gels stained with picrosirius red stain (pink color), untreated (leftmost) and treated with increasing concentration of 50 μM, 100 μM and 200 μM (left to right) of methylglyoxal (top row), glyoxal (middle row) and 3-deoxyglucosone (bottom row) for 1 d. (B) Photomicrographs of polymerized Collagen I gels stained with picrosirius red stain (pink color), untreated (leftmost) and treated with increasing concentration of 50 μM, 100 μM and 200 μM (left to right) of methylglyoxal (top row), glyoxal (middle row) and 3-deoxyglucosone (bottom row) for 4 d. Experiment carried out for more than three biological replicates. Scale bar for (A) and (B) = 1 cm. Three or more independent repeats were performed for each experiment.

### MGO and GO lead to denaturation of secondary structure of Collagen I

We sought to confirm increased crosslinking of Collagen I that we observed using picrosirius staining. Circular Dichroism confirmed whether there is any denaturation of the secondary structure of Collagen I. Pre-polymerized Collagen I samples of concentration 0.1 mg/mL were treated with 1X PBS (control) and with 200 μM of MGO and GO for 4 d. Untreated control Collagen I showed peaks at 197 nm and 222 nm wavelength. A reduction in the molar ellipticity was observed at 197 nm wavelength in case of both MGO- and GO-treated Collagen I, indicating denaturation of the secondary structure of Collagen I in the presence of these two glycating agents (Supplementary Figure 2).

Increased crosslinking of Collagen I that disrupts its secondary structure decreases the availability of receptors for cell adhesion leading to a significant decrease in the number of adherent cells. The cells that adhere do not form their characteristic spindle shape, which further causes a reduction in the contractile activity of the cells and hence, reduction in the velocity and accumulated distance of the cells. This complements the adhesion and migration results we obtain for endothelial cells and cancer cells in our experiments.

## DISCUSSION

Three reactive dicarbonyls, MGO, GO, and 3-DG, have been investigated widely in context of Type 2 diabetes mellitus and aging; most studies have focused on their direct impact on the homeostasis of cells and the organization of tissues and organs^7,15,18–20,26^. Our study builds on previous observations made by our group and others on the effect of dicarbonyls on the extracellular matrix which serves as the scaffolding of tissues as well as reinforces the homeostatic state of their parenchyma^7,15,18,26^; it then goes further and uses these assays to compare the specific effects of individual dicarbonyls on tissue organization and cellular morphomigratory traits. The results obtained for both cancerous and non-cancerous cells suggest a clear perturbation of both migration and adhesion on matrices treated with MGO at even low concentrations and with GO at high concentrations. 3-DG and low concentrations of GO seem relatively ineffectual at perturbing cell-matrix interactions. The alteration in adhesion on matrices treated with MGO and GO is phenomenologically similar for both endothelia and breast cancer cells, but the implications are drastically different. Untransformed cells upon loss in adhesion undergo anoikis and leave behind tissues that are vulnerable to invasion by pathological cell populations such as in cancer metastasis^38–40^. Cancer cells, on the other hand, are programmed to survive when detached from matrices; detachment, in fact, aids their invasive progression^40–48^.

We observe higher migration of cancer cells on C2-α-dicarbonyl-treated lrBM: the reason is the dissolution of the matrix leading to a likely measurement of cell migration on the underlying plastic substrata: lrBM is a confining sheet-like barrier for cancer cells and has to be breached for the cancer cells to migrate into stroma ^49,50^. In contrast, we observe a decrease in migration of cancer cells on Collagen I treated with MGO and high levels of GO: although this may suggest an impaired kinetics of progression mediated by dicarbonyls, the association is worth re-examining, when the assay is made to further approximate in vivo histological complexity by incorporating normal untransformed parenchymal or stromal cells around the cancer cells. In this regard, the decreased migration of endothelia on dicarbonyl-treated matrix suggests impaired vasculogenesis and wound healing under carbonyl stress^51–53^. Lower migration of cells on glycated Collagen I substrata is associated with a decreased entropy of elongation, which implies that the cells are intrinsically less deformable and more rigid in such milieu. This suggests that mechanical cues from the glycated matrices are interpreted in distinct ways from control substrata by migrating cells resulting in altered cortical organization of the cellular cytoskeleton. We aim to test this hypothesis in future experiments through a combination of time lapse imaging and traction force microscopy.

The migration and adhesion data for both Collagen I and lrBM suggested a change in the matrix properties on exposure to dicarbonyl stressors. We observe dissolution of lrBM upon treatment with MGO and GO, with a more profound impact with MGO than similar concentrations of GO. There is no such change observed in case of treatment with 3-DG. LrBM, on the hand, dissolves in the presence of MGO and GO, faster under treatment with MGO than GO. In contrast, we observe insignificant dissolution in 3-DG-treated lrBM. The existent literature reports a similar chemical reactivity order of dicarbonyl stressors with GSH, where the difference in hydration of the three stressors has been considered to be the reason behind their order of reactivity^19^. The keto group of MGO is less susceptible to hydration than the two aldehyde groups of GO, making the former more electrophilic, and reactive towards nucleophilic proteins^54,55^. Additionally, 3-DG forms a cyclic structure on hydration which contributes to its relatively lowest reactivity^55^. These findings imply that C2-α-dicarbonyls may not just be more reactive to extracellular matrices like Collagen I and lrBM than C6-α-dicarbonyls, there is also difference in the behavior of two different C2-α-dicarbonyls in their reaction with ECMs. However, the reactivity of MGO makes it also susceptible to glyoxalase-based conversion to D-lactic acid within cells^19,56,57^. In the extracellular milieu, the lack of dicarbonyl-processing enzymes such as glyoxalase, aldoketoreductase and aldehydyde dehydrogenase, all of which are known to act on and decrease levels of dicarbonyls, results in MGO exerting a far greater pathological effect on matrix structure and organization^4,7,15,18^.

We wish to highlight the limitations of our study. While our experiments have been carried out with a triple negative breast cancer cell line and an endothelial cell line, the results may vary qualitatively or quantitatively for cancer cells originating from other tissues as well as non-cancerous cells. We have restricted our comparative analysis to three widely used dicarbonyls, but would like to extend the same to other dicarbonyls such as 3-DGal (3-Deoxygalactosone) and diacetyls which have their distinct signatures of cytotoxicity but whose effects on matrices are poorly, if at all, understood^20^. Such limitations notwithstanding, we aim to utilize the results obtained from this study to focus on quenching specific dicarbonyl stressors whose accumulation potentiate organopathies through attenuating the homeostatic functions of extracellular matrices.

## MATERIALS AND METHODS

### Cell culture

This study utilizes two cell lines, triple negative breast cancer cell line, MDA-MB-231 and endothelial cell line, TeloHAEC. MDA-MB-231 (ATCC) were prepared to stably express red fluorescent protein (RFP) using lentiviral transduction. MDA-MB-231 RFP cells were cultured in DMEM F-12 (HiMedia, AT007F) supplemented with 10% fetal bovine serum (Gibco, 10270). For this study, they have been trypsinized at a dilution of 1:4 with 0.25% trypsin and 0.02% EDTA (HiMedia, TCL007). TeloHAEC (ATCC) were prepared to stably express green fluorescent protein (GFP) using lentiviral transduction. TeloHAEC GFP cells were cultured in EBM-2 Media (Lonza, #CC3302). Similar to MDA-MB-231 RFP, they have been trypsinized at a dilution of 1:4 with 0.25% trypsin and 0.02% EDTA (HiMedia, TCL007).

### Adhesion of cancer cells and endothelial cells on lrBM

Laminin-rich basement membrane coats were prepared by directly adding 100 μL of Matrigel™, consisting of Collagen IV, Laminin, Entactin, and Perlecan, of the stock concentration 9.8 mg/mL in the form of a coat on a chamber well and were allowed to solidify at 37°C for 30 min. The coats were incubated at 37°C in 1X PBS (untreated control) and in 50 μM, 100 μM and 200 μM of methylglyoxal, glyoxal and 3-deoxyglucosone for 1 d. After incubation, the treatment was removed, the coats were properly washed with 1X PBS to remove any residual unreacted glycating agents. Breast cancer cells MDA-MB-231 and endothelial cells TeloHAEC were added to the wells at a density of 10,000 cells per well and were allowed to adhere to the matrix for 30 min inside an incubator at 37°C and at 5% CO_2_ conditions. The cell suspension was then removed and adhered cells were fixed using 4% formaldehyde for 30 min at 4 °C. Cells were imaged using Olympus IX83 inverted fluorescence microscope fitted with Aurox structured illumination spinning disk setup at 10X magnification. Images were processed in open-source platform Fiji.^58^ Cells were counted using ImageJ software and graphs were plotted for three biological replicates with 10 fields of view per replicate.

### Migration of endothelial cells on lrBM

100 μL coats of lrBM were prepared and solidified at 37°C for 30 min. The coats were incubated in 37°C with 1X PBS (control) and 50 μM of methylglyoxal, glyoxal and 3-deoxyglucosone for 1 d. The coats were washed properly after 1 d with 1X PBS and endothelial cells TeloHAEC were added to the wells at a density of 10,000 cells per well and kept for 24 h to allow the cells to attach to lrBM. Images were taken in inverted epifluorescence microscope (IX83 Olympus, Tokyo, Japan) at 4× objective. Images were processed and converted to RGB color. Angiogenesis Analyzer plugin for ImageJ was used to evaluate the endothelial tube formation that occurs when endothelial cells are added to lrBM, where they form meshed pseudo-capillary networks^59^. The angiogenesis plugin analyser settings were set based on the behavioural properties of TeloHAEC with minimum object size as 10 pixels, minimum branch size ad 25 pixels. Artifactual loop size as 850 pixels, isolated element size threshold to be around 25 pixels, and master segment size as 30 pixels. Three iterations are run for the plugin and the final output shows the isolated elements marked in blue color while the main branches marked in green (Supplementary Figure 3). The number of isolated elements was quantified and plotted on a graph for three biological replicates with 28 fields in total.

### Migration of cancer cells on lrBM

100 μL lrBM coats were prepared on an eight well chamber and were allowed to solidify at a temperature of 37°C for 30 min. The coats were incubated in 37°C with 1X PBS (untreated) and with 50 μM methylglyoxal, glyoxal and 3-deoxyglucosone treatment for 1 d. After 1 d, the treatment was removed, the coats were properly washed with 1X PBS to remove any residual unreacted glycating agents. Breast cancer cells MDA-MB-231 were added to the wells at a density of 10,000 cells per well and kept overnight to allow the cells to attach to lrBM. Time lapse imaging in inverted epifluorescence microscope (IX83 Olympus, Tokyo, Japan) at 20× objective was then carried out for 3 h, with snapshots taken at every 2 min interval. Images were processed in open source platform Fiji^58^. Morphomigratory parameters of mean velocity, accumulated distance, elongation and elongation entropy were then assessed using a metric toolkit that measures the trajectories of migration using Shannon entropic distribution^36^. The results obtained were plotted for three biological replicates with 50 cells per replicate for cancer cells and 25 cells per replicate for endothelial cells.

### Alcian Blue staining of lrBM

LrBM gels of 50 μL were prepared and solidified at 37°C for 30 min. They were treated with 1X PBS and increasing concentrations of 50 μM, 100 μM and 200 μM of MGO, GO and 3-DG for 1 d and 4 d. After their treatment duration was over, the gels were properly washed with 1X PBS and fixed with 1% formaldehyde at 37°C for 30 min. Alcian blue stain was prepared by adding 1% w/V Alcian blue 8GX (HiMedia, TC359-10G) in 3% acetic acid solution at pH 2.5. The gels were treated with 3% acetic acid solution for 3 min, followed by staining with Alcian blue for 30 min at 37°C. The gels were washed with 1X PBS and imaged in a dark room with a uniform light source from the bottom of the plate using a Nikon Z6 III camera.

### Adhesion of cancer cells and endothelial cells on Collagen I

Collagen I (Gibco, A1048301) of stock concentration of 3 mg/mL was diluted to a working concentration of 1 mg/mL after neutralizing it using 2N Sodium Hydroxide (NaOH), 10X DMEM, Dulbecco’s Modified Eagle Medium (HiMedia, AT006F) and 1X PBS (Phosphate buffered saline). 100 μL coats of Collagen I were prepared on an eight well chamber and were allowed to polymerize at a temperature of 37°C for 30 min. The coats were incubated in 37°C with 1X PBS (untreated control) and with 50 μM, 100 μM and 200 μM of methylglyoxal, glyoxal and 3-deoxyglucosone for 4 d. MGO, GO and 3-DG were replenished after 2 d. After 4 d, the treatment was removed, the coats were properly washed with 1X PBS to remove any residual unreacted glycating agents. Breast cancer cells MDA-MB-231 and endothelial cells TeloHAEC were added to the wells at a density of 10,000 cells per well and were allowed to adhere to the matrix inside an incubator at 37 °C and at 5% CO_2_ conditions. The cell suspension was removed after an hour of incubation and adhered cells were fixed using 4% formaldehyde for 30 min at 4°C. Cells were then imaged using Olympus IX83 inverted fluorescence microscope fitted with Aurox structured illumination spinning disk setup at 10X magnification. Images were processed, cells were counted using ImageJ software and graphs were plotted for three biological replicates with 10 fields of view per replicate.

### Migration of cancer cells and endothelial cells on Collagen I

100 μL coat of Collagen I was prepared on an eight well chamber and were allowed to polymerize at a temperature of 37°C for 30 min. The coats were incubated in 37°C with 1X PBS (untreated control) and 200 μM of methylglyoxal, glyoxal and 3-deoxyglucosone for 4 d. MGO, GO and 3-DG were replenished after 2 d. After 4 d, the treatment was removed, the coats were properly washed with 1X PBS to remove any residual unreacted glycating agents. Breast cancer cells MDA-MB-231 and endothelial cells TeloHAEC were added to the wells at a density of 10,000 cells per well and kept overnight to allow the cells to attach to Collagen I. Time lapse imaging in inverted epifluorescence microscope (IX83 Olympus, Tokyo, Japan) at 20× objective was then carried out for 3 h, with snapshots taken at every 2 min interval. Images were processed, morphomigratory parameters of mean velocity, accumulated distance, elongation and elongation entropy were assessed and graphs were plotted for three biological replicates with 50 cells per replicate.

### Picrosirius staining of Collagen I

Collagen I spherical gels of 50 μL were prepared and allowed to polymerize at 37°C for 30 min. They were then treated with 1X PBS and increasing concentrations of 50 μM, 100 μM and 200 μM of MGO, GO and 3-DG for 1 d and 4 d. After 1 d and 4 d respectively, the treatment was removed and the gels were properly washed with 1X PBS and distilled water to remove any residual unattached MGO, GO and 3-DG. 10 mL Picric acid solution was prepared by heating 0.12 g of Picric acid and 10 mL distilled water on hotplate for 5-10 min after mixing them properly. 0.01 g of Direct Red 80 (Sigma–Aldrich, 365548-5G) was then added to the mix to prepare a homogenous solution of picric acid. This solution was then added to the gels and incubated for 2 h at room temperature. The gels were washed with 0.05% acetic acid, absolute alcohol, and 1X PBS. They were stored in 1X PBS for 24 h to get rid of the excess stain and imaged in a dark room with a uniform light source from the bottom of the plate using a Nikon Z6 III camera.

### Circular dichroism of pre-polymerized Collagen I

Collagen I was diluted to a concentration of 0.1mg/mL in 0.1% acetic acid. 500 μL solution consisting of 0.1 mg/mL Collagen I untreated (1X PBS) and treated with 200 μM of methylglyoxal and glyoxal was prepared and kept for 4 d at 4°C away from light. The samples were taken for circular dichroism spectroscopy, performed using Jasco-815 spectropolarimeter purged with inert gas in the spectral range of 190 nm-260 nm with the help of a rectangular quartz cuvette that has an optical path of 0.1 cm. A graph of molar ellipticity (in degrees cm2 dmol-1) versus wavelength (in nm) was plotted for the spectral range of 190 nm – 260 nm.

## ACKNOWLEDGEMENT

This work was supported by the Wellcome Trust/DBT India Alliance Fellowship/Grant [IA/I/17/2/503312] awarded to R.B. It was also supported by the Indo-French Centre for the Promotion of Advanced Research (69T08-2) to R.B. BS acknowledges support from the Prime Ministers Research Fellowship. This study acknowledges funding from the Longevity India Initiative supported from the Prashanth Prakash Family Foundation. The authors thank MeITY, Government of India. They thank Neeru Yadav, Imnatoshi Jamir and Allu Sainandan Reddy for their assistance with Nikon Z6 III camera and C. Inbasekar for their assistance with circular dichroism that helped improve the manuscript. The opinions expressed in this paper are those of the authors and not those of the Prashanth Prakash Family Foundation.

## DATA AVAILABILITY STATEMENT

The data that support the findings of this study are available from the corresponding author upon reasonable request.

