## Supplementary data for "C2-α-dicarbonyls selectively impair morphodynamic interactions between cells and extracellular matrix"

A.

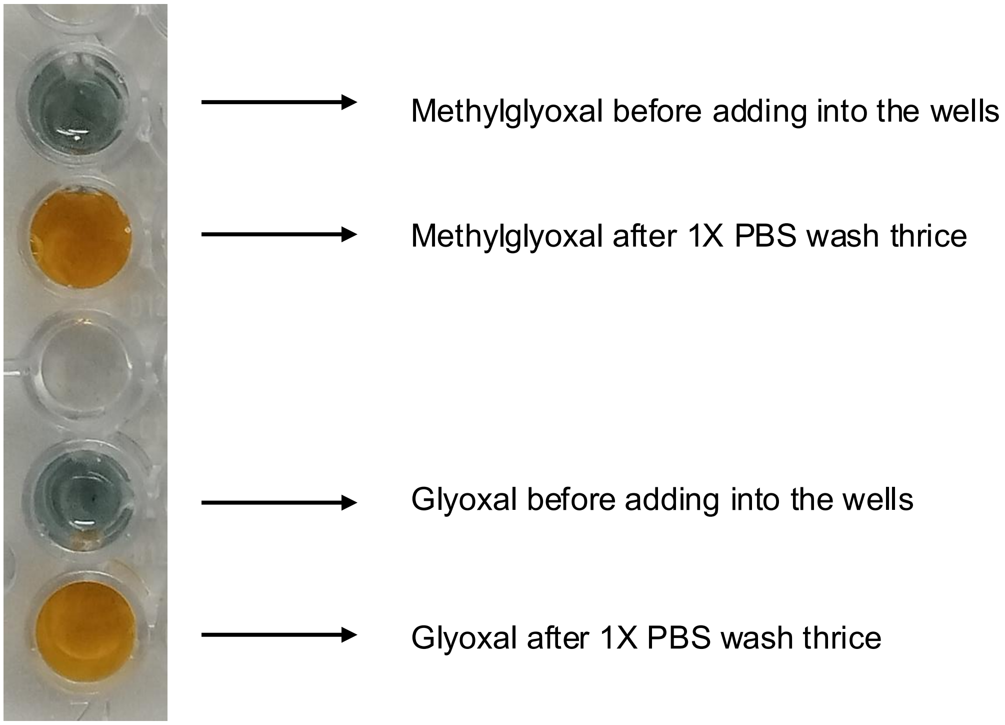


B.

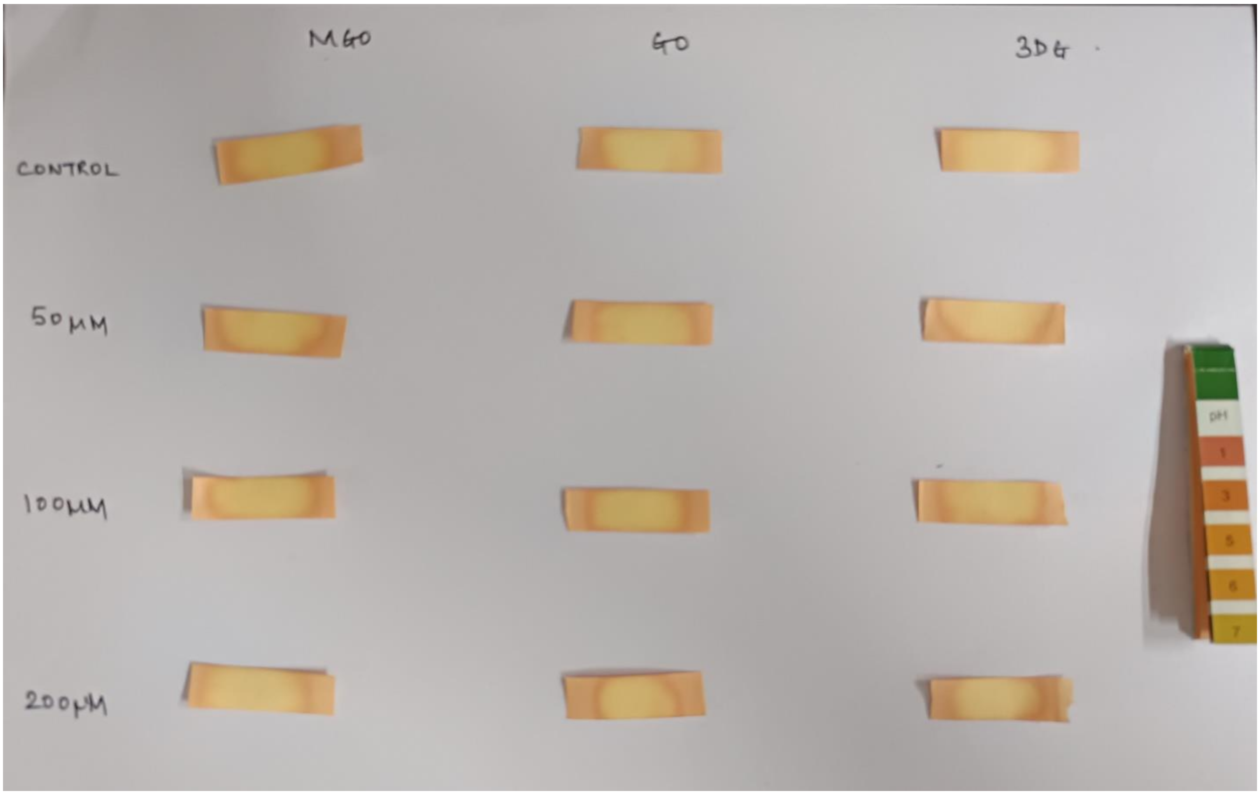


Supplementary Figure 1: **Confirmatory tests for the presence and wash of dicarbonyl stressors** (A) Photomicrographs of chromic acid staining of methylglyoxal before addition into the wells, methylglyoxal after a proper 1X PBS wash, glyoxal before addition into the wells and glyoxal after a proper 1X PBS wash (top to bottom) (B) Photomicrographs of control (1X PBS) and 50 µM, 100 µM and 200 µM (top to bottom) of methylglyoxal (left), glyoxal (middle) and 3-deoxyglucosone (right) deposited on pH strips. pH label (rightmost) to indicate approximate pH value obtained for the samples.

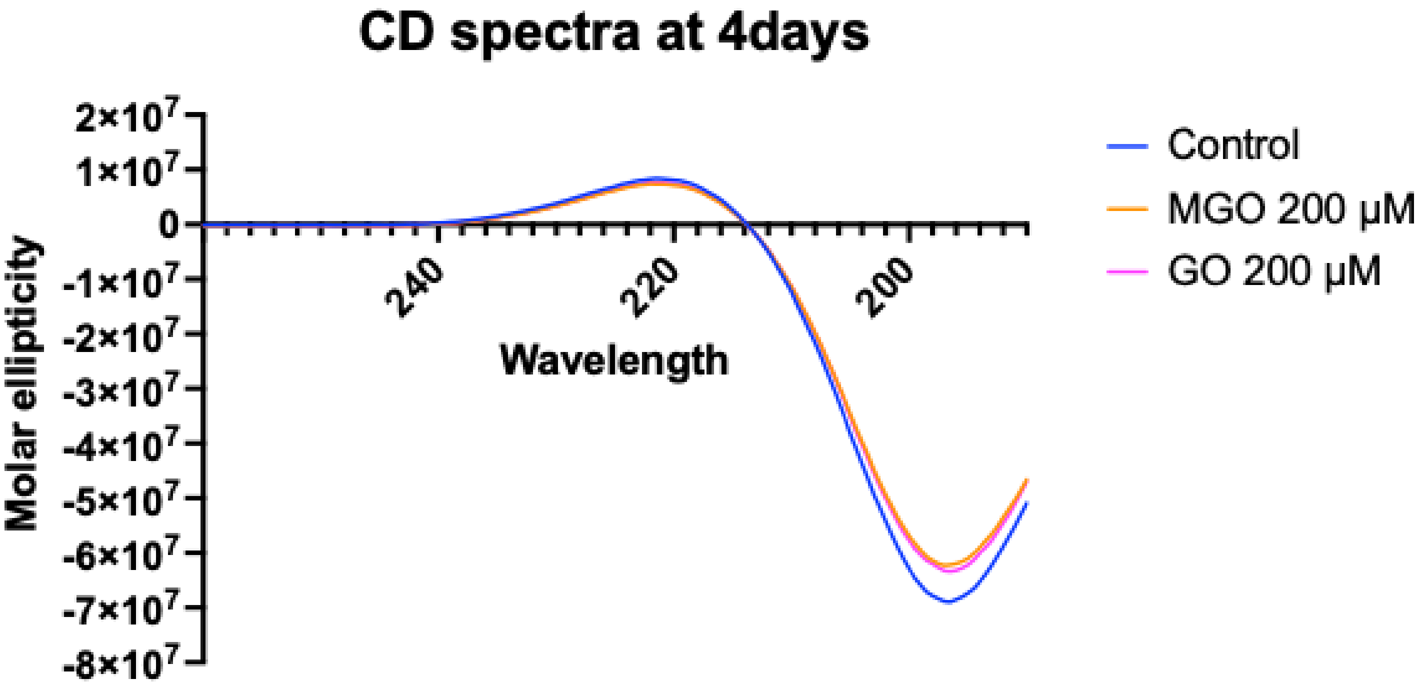


Supplementary Figure 2: **MGO and GO denature the secondary structure of Collagen I,** XY graph plots of Molar ellipticity (degrees $cm^2$ $dmol^{-1}$) versus Wavelength (nm) obtained for Collagen I treated with 1X PBS and 200 μM of methylglyoxal and glyoxal for 4 d, measured using Circular Dichroism. Data collected for three biological replicates.

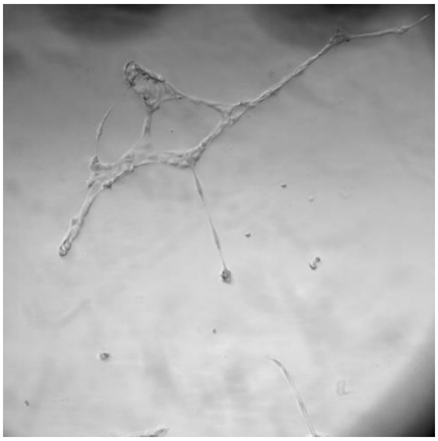

1. Open file in ImageJ and convert into RGB colour.

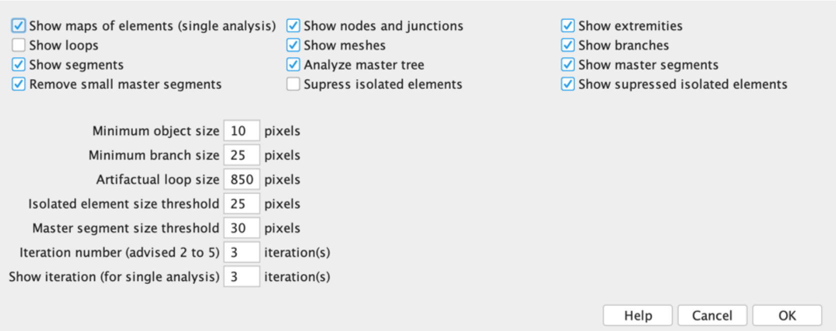


2. Open Angiogensis Analyser plugin for Phase constrast imagine and set parameters as shown.

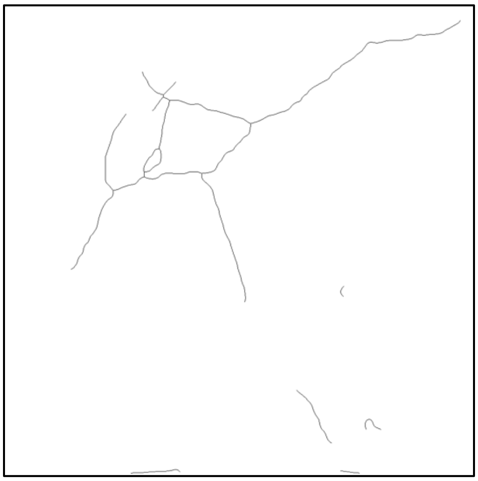

3. Map of the final tree is generated.

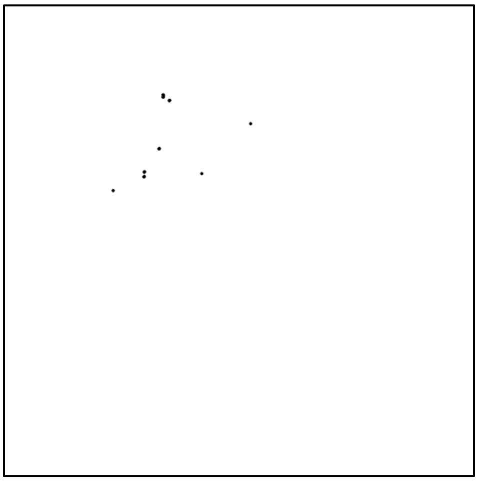

4. Map of the nodes is generated.

| | A | B | C | D | E | |
|---|---|---|---|---|---|---|
| 1 | Image Nar | Analysed a | Nb extrem | Nb nodes | Nb Junctic | Nb |
| 2 | 004-tr | 1048576 | 17 | 21 | 8 | |

5. Statistics table is generated.

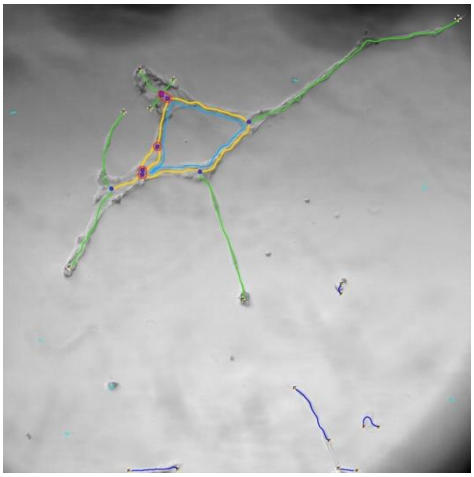

6. Final tree representing the mesh formed by the endothelial cells on IrBM is generated.

Supplementary Figure 3: **Protocol for Angiogenesis Assay,** Schematic Representation of the protocol of endothelial tube formation assay for TeloHAEC cell line using Angiogenesis ImageJ plugin.
